# A system for generating reversible knockout cell lines

**DOI:** 10.1101/2022.05.25.493512

**Authors:** Kacee A. DiTacchio, Luciano DiTacchio

**Affiliations:** Synexin, PO BOX 485, Carlsbad CA 92018

## Abstract

A powerful use of gene editing technology is the generation of genetic-null (knockout) mutants of human immortalized cell lines. However, a problem that arises in the process of generating such edited lines is that to ensure edit homogeneity, clonal selection must often be performed and, given the genomic instability of immortalized cell lines and the high division number inherent to the clonal selection process, the resulting clone can exhibit genotypic and phenotypic differences from the parental line. Here, we present a system that allows for the generation of genetic null cell lines with an excisable cassette. This system allows for the generation of suitable controls without the need of further rounds of clonal selection.

## Results

Cas9/CRISPR-based gene editing technology has revolutionized biomedical research (Hryhorowicz et al., 2017; Ma et al., 2014; Zhang et al., 2014). A powerful use of this technology is the generation of genetic-null (knockout) mutants of human immortalized cell lines, something that was previously technologically not possible. There are two main approaches to generate such knockout cell lines, both of which rely on endogenous DNA repair mechanisms (Zhang et al., 2016). In one of these approaches, Cas9-induced breaks in DNA are used to trigger the error-prone non-homologous end joining (NHEJ) repair, which leads to insertion-deletion (indel) mutations at the repair junction. Although high in frequency, these indel mutations do not always result in the frameshifts required for disruption of gene expression and, consequentially, clonal selection is frequently required to ensure abrogation of gene expression has been achieved (Guo et al., 2018). A separate approach exploits the homologous DNA recombination (HDR) repair mechanism by providing an DNA template harboring desired modifications flanked by unaltered sequences. Although this approach ensures that the targeted allele is modified as desired, its efficiency is low and, as with indel-based editing, clonal identification and selection must be carried out.

A key hurdle for both indel-mediated and HDR-mediated face relates to the lack of proper wild-type or unmodified controls. Frequently, what is used as a control is either the parental line, clones obtained from a “mock” modification process, or a combination of the two. Yet, the genomic instability of immortalized lines leads to significant drift in the characteristics of the resulting clones compared with the parental line (Barnes et al., 2006; Tharmalingam et al., 2018).

We have established a novel gene editing approach that allows us to disrupt gene expression and that can be reversed in an expanded clone population without the need for further selection –thus generating as ideal a control as currently technically possible. In our approach, we use homologous recombination to split an exon and introduce a selectable and excisable disruption cassette that terminates both transcription and translation (Figure. 1). Following selection of edited cells, the disruption cassette can then be excised by the action of Cre recombinase, rescuing gene expression.

**Figure 1.**
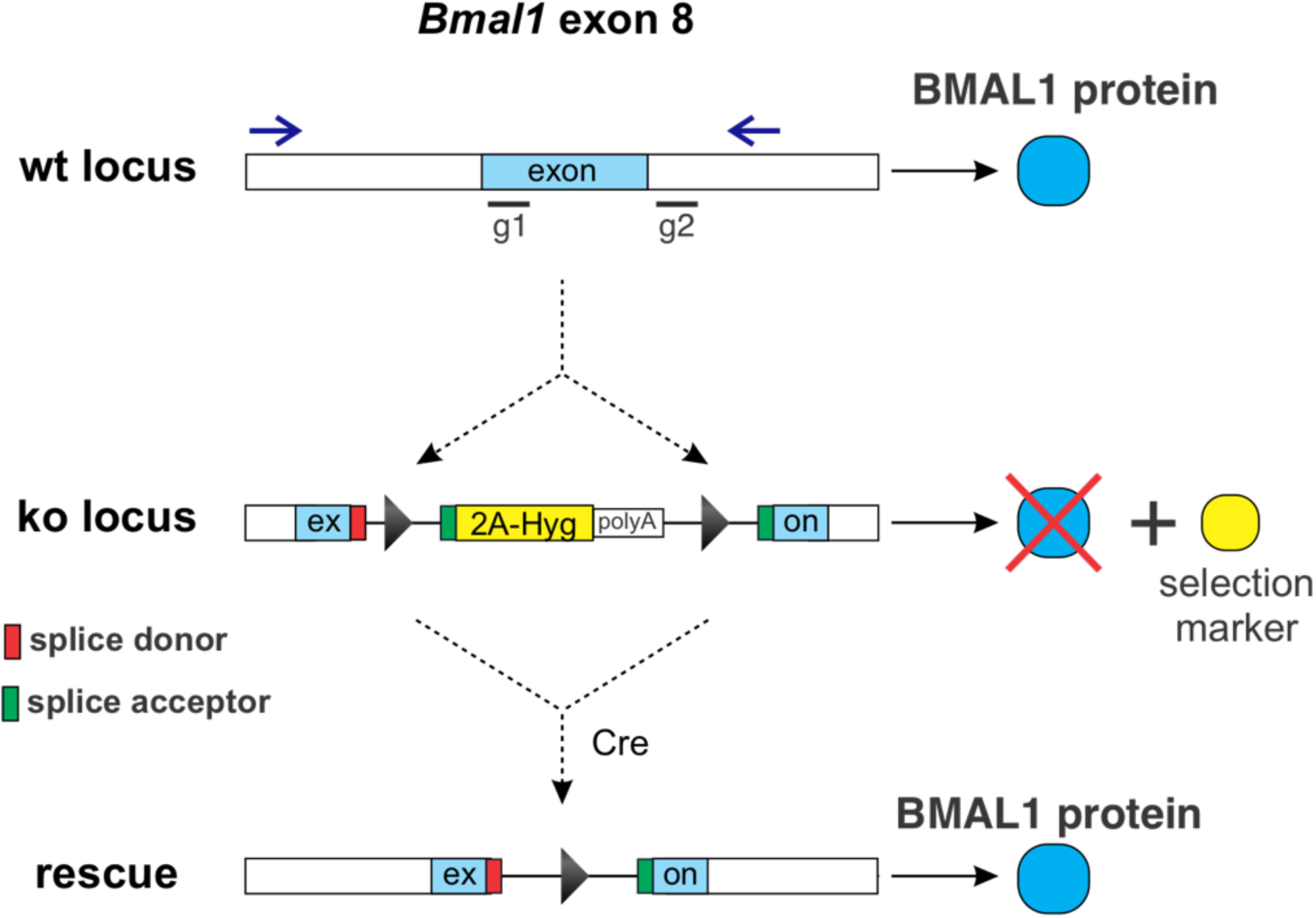
Exon-splitting approach used to ablate gene expression. Blue arrows denote the location of primers used for genotyping. Black bars labeled g1 and g2 are the location of gRNA target sequences. The ablation cassette diverts splicing of the endogenous transcript to generate a truncated BMAL1 protein and a selection marker, while forcing transcription termination. Upon identification and expansion of a knockout clone, the disruption allele is excised by expression of Cre recombinase.

As proof of principle, we applied our system to abrogate expression of the key circadian clock component BMAL1 (ARNTL) in U2OS cells given that *Bmal1* loss in this line has been shown to have robust phenotypes (Baggs et al., 2009; Vollmers et al., 2008). First, we introduced a cassette into exon 8 of the *Bmal1* gene, which codes for the basic Helix-Loop-Helix DNA-binding domain of BMAL1 and whose targeting has been used to generate genetic-null animals (Storch et al., 2007). Following antibiotic selection, single cells were plated in 96-well plates and three clonal lines were identified via PCR genotyping as both harboring the modified allele and lacking an unmodified one (Figure 2). We observed loss of BMAL1 expression, and the expected associated loss of circadian rhythmicity as assessed by real-time luminescence measurements of a Bmal1 promoter-driven Luciferase reporter construct. We were able to rescue both BMAL1 expression and circadian rhythmicity by expression of Cre recombinase, which we delivered into the expanded clone population via an adenovirus vector (Figure 3). Of note, even though rhythmicity is restored in cells where BMAL1 expression has been rescued, the circadian profile is not identical to the original cell line (Figure 4), which is consistent with the aforementioned changes that occur during the clone selection process.

**Figure 2.**
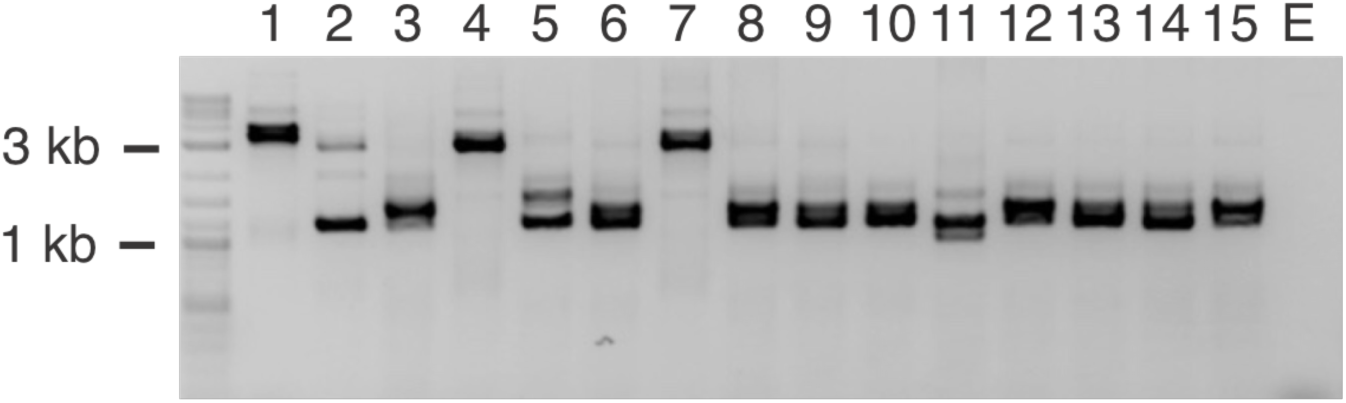
Identification of U2OS Bmal1 knockout clones by PCR genotyping. The unmodified allele yields a 1372 bp amplicon, whereas presence of the disruption cassette yields one of ∼3200 bp in length. E, no template control.

**Figure 3.**
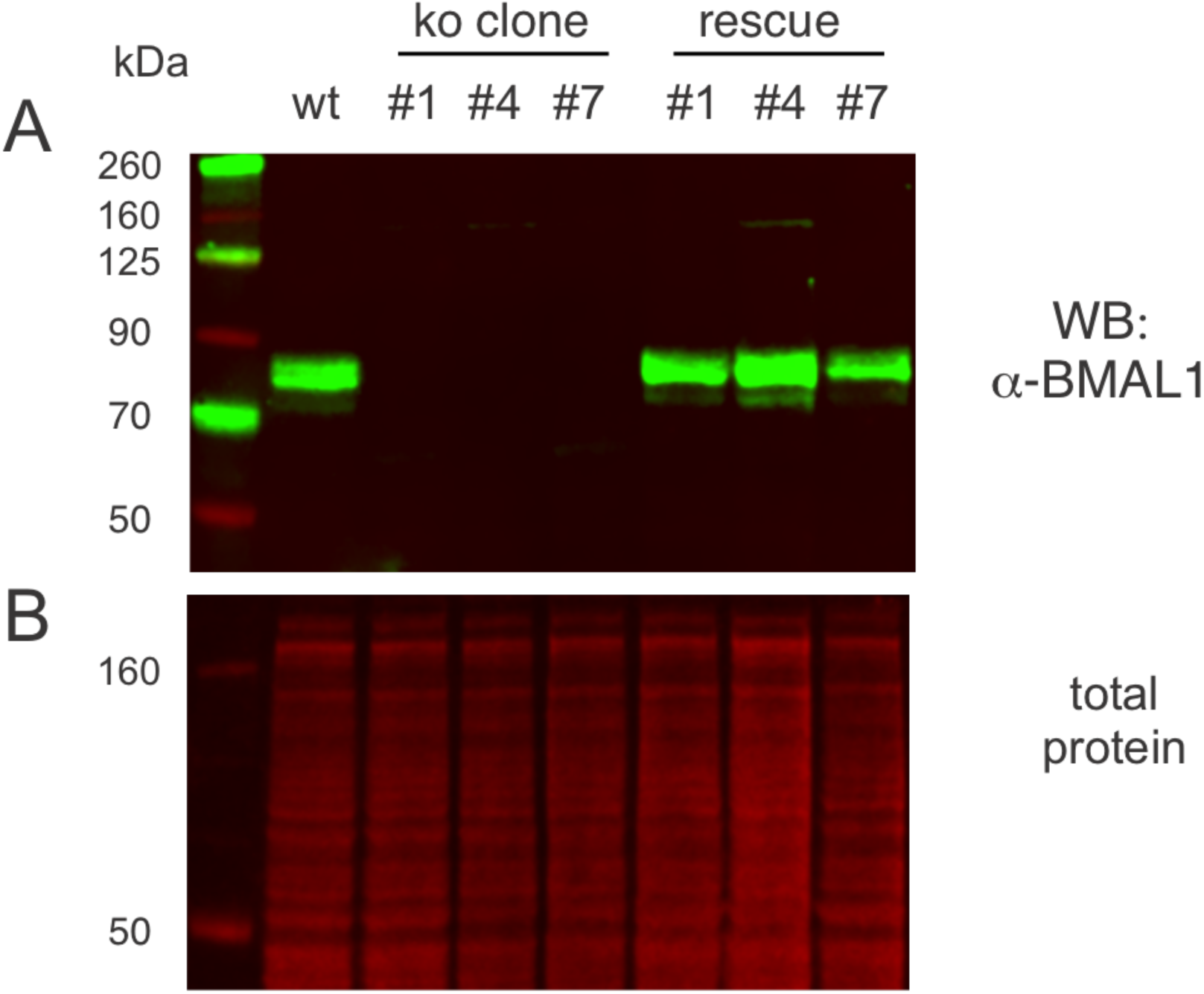
Disruption of the Bmal1 locus in U2OS cells by insertion of a reversible cassette and its rescue by induced recombination. A) Anti-BMAL1 immunoblot of extracts derived from the parental U2OS line (wt), three knockout clones (#1, #4, #7), and their rescued controls. B) Corresponding total protein stain for the blot shown.

**Figure 4.**
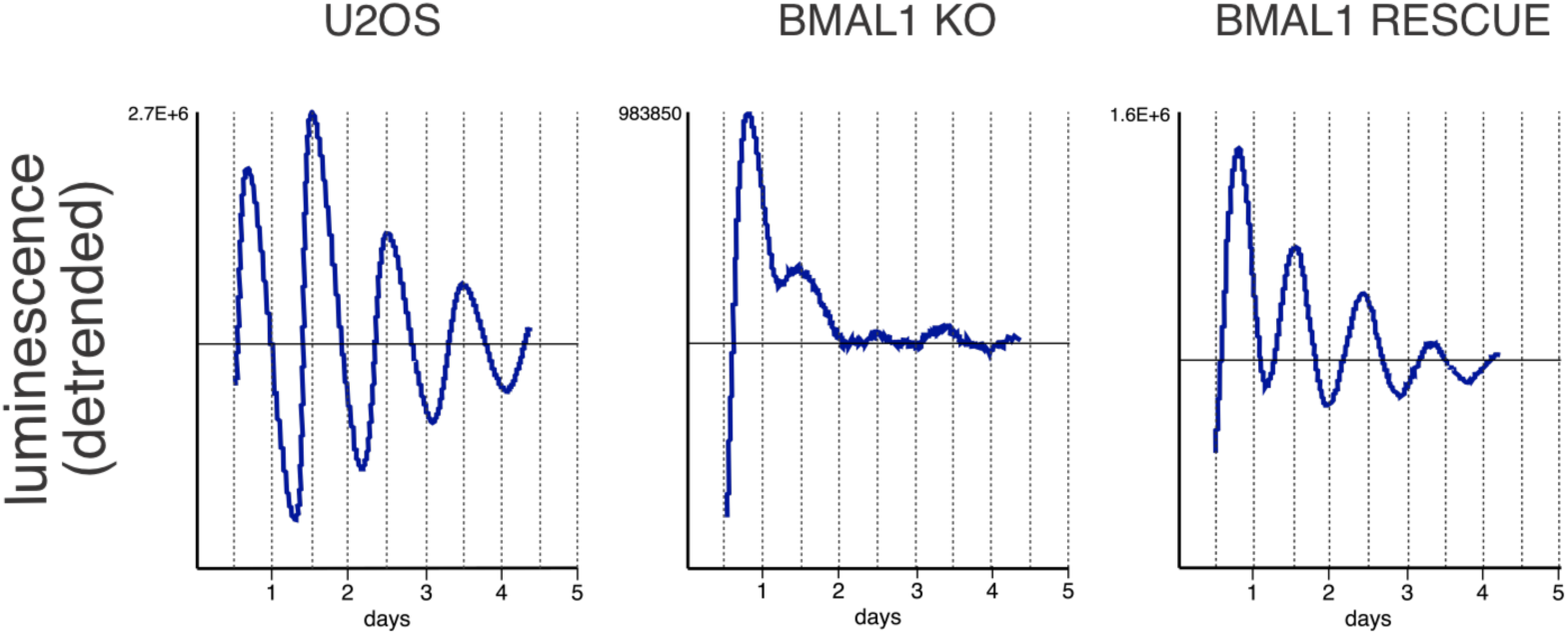
Real-time luciferase measurements show that disruption of the *Bmal1* locus in U2OS cells by insertion of a reversible cassette leads to loss of circadian rhythms, and their rescue upon reversal to the WT. U2OS cells, U2OS BMAL1^KO^ cells (clone #4), and the corresponding rescue control cells (rescue control #4) were transfected with a Bmal1-luciferase reporter construct and luminescence measured continuously for 5 days.

## Materials and Methods

### Cell Culture, Transfections, and Luciferase Assays

U2OS cells were cultured in Dulbecco’s Modified Eagle Medium (DMEM) (Corning Cat# 10-013-CV) supplemented with 10% FBS (Atlanta Biologicals Cat# S11595H) and 1% antibiotic/antimycotics (Thermo Fisher Cat# 15240062) in a 37 °C incubator at 5% CO_2_.

Transfections were performed using Trans-IT LT1 (TLT-1) (Mirus Bio) according to the manufacturer’s instruction.

Gene editing was performed by transfecting U2OS cells with plasmids for Cas9 and gRNAs in the presence of a donor template consisting of a disruption cassette flanked by sequences homologous to the endogenous Bmal1 exon 8 locus. Four days after transfection, single cells were seeded onto 96 well tissue culture treated plates and kept under antibiotic selection and following identification by genotyping, knockout clones were expanded.

To excise the disruption cassette, we seeded 2 million cells of the desired clone onto a 10 cm dish and delivered Cre via adenovirus (10 x106 pfu per 2 million cells, Vector BioLabs).

Real-time luciferase assays were performed in a 96-well plate format by reverse transfecting cells with 300 ng *Bmal1*-Luciferase reporter construct and measuring luminescence in a temperature-controlled microplate reader for up to 7 days.

### Immunoblotting

Whole-cell lysates from cells were prepared using lysis buffer (150 mM NaCl, 50 mM Tris-HCl, 0.5% TX-100, 0.5% NP-40, 0.25% Sodium Deoxycholate 0.025% SDS) supplemented with EDTA-free protease inhibitor cocktail and phosphatase inhibitors (Roche). Lysates were clarified by centrifugation, resolved by SDS-PAGE, transferred onto a PVDF membrane, and probed with an anti-BMAL1 antibody (Abcam #3350).

Immunoblotting was performed with an Odyssey Imaging System (LiCor) and related reagents. Total protein content was visualized with Revert reagent (LiCor).

### Oligonucleotides

All oligonucleotides were ordered from IDT.

Bmal1 genotyping Fwd: GTCCTCAGTTCTTCTGCCGTT

Bmal1 genotyoing Rev: GCACCTACATTTCTGCGTGT

Guide RNAs targeting the Bmal1 exon 8 region

g1: CTCACAGTCAGATTGAAAAG

g2: GATAGGAACATTCTAGGCAA

